# Microbiome of the Black Sea water column analyzed by genome centric metagenomics

**DOI:** 10.1101/2020.10.30.362129

**Authors:** Pedro J. Cabello-Yeves, Cristiana Callieri, Antonio Picazo, Maliheh Mehrshad, Jose M. Haro-Moreno, Juan J. Roda-Garcia, Nina Dzhembekova, Violeta Slabakova, Nataliya Slabakova, Snejana Moncheva, Francisco Rodriguez-Valera

## Abstract

**Background:** The Black Sea is the largest brackish water body in the world, although it is connected to the Mediterranean Sea and presents an upper water layer similar to some regions of the former albeit with lower salinity and (mostly) temperature. In spite of its well-known hydrology and physico chemistry, this enormous water mass remains poorly studied at the microbial genomics level.

**Results:** We have sampled its different water masses and analyzed the microbiome by classic and genome-resolved metagenomics generating a large number of metagenome-assembled genomes (MAGs) from them. The oxic zone presents many similarities to the global ocean while the euxinic water mass has similarities to other similar aquatic environments of marine or freshwater (meromictic monimolimnion strata) origin. The MAG collection represents very well the different types of metabolisms expected in this kind of environments and includes Cyanobacteria (*Synechococcus*), photoheterotrophs (largely with marine relatives), facultative/microaerophilic microbes again largely marine, chemolithotrophs (N and S oxidizers) and a large number of anaerobes, mostly sulfate reducers but also a few methanogens and a large number of “dark matter” streamlined genomes of largely unpredictable ecology.

**Conclusions:** The Black Sea presents a mixture of similarities to other water bodies. The photic zone has many microbes in common with that of the Mediterranean with the relevant exception of the absence of *Prochlorococcus*. The chemocline already presents very different characteristics with many examples of chemolithotrophic metabolism (*Thioglobus*) and facultatively anaerobic microbes. Finally the euxinic anaerobic zone presents, as expected, features in common with the bottom of meromictic lakes with a massive dominance of sulfate reduction as energy generating metabolism and a small but detectable methanogenesis.We are adding critical information about this unique and important ecosystem and its microbiome.

## Background

The Black Sea is the inner arm of the Mediterranean basin. Nearly severed from the rest by the tectonic movement of the African plate, it is only connected to the rest of the Mediterranean Sea by the narrow but deep strain of the Bosporus. The Black Sea has a positive hydric balance i.e. receives more freshwater than lost by evaporation and hence contains less salt (from 0.73 % in epipelagic to 2.2 % in meso-bathypelagic waters) than the Mediterranean (3.8 %) proper. In addition, the large watershed and riverine inputs lead to a richer nutrient status (meso-eutrophic) and permanent stratification with a colder, more saline deep water mass that remains anaerobic and largely euxinic below 150-200 m [1–3]. All these properties make the Black Sea a unique brackish-marine environment. Its great depth (average depth 1253 m with a maximum of 2212 m) makes this system much more stable than other brackish inland water bodies like the Baltic Sea in which the anaerobic compartment is only a recent development due to anthropic impact [4].

Although a few studies have been carried out by metagenomics and metagenome-assembled genomes (MAGs) reconstruction, the information available in databases about this unique environment is scarce. A recent study showed for the first time the microbial structure of the sulfidic waters of 1000 m depth [5], mainly dominated by sulfate reducers (Desulfobacterota and Chloroflexi classes such as Dehalococcoidia or Anaerolinea), associated DOM degraders (Marinimicrobia, Cloacimonetes) and streamlined uncultured taxa such as Omnitrophica, Parcubacteria or Woesearchaeota. At the genome level, Black Sea microbes remain largely unknown. Only 10 MAGs have been studied from the abovementioned study [5], 179 MAGs from 50-2000 m have been recently deposited into Genbank (PRJNA649215) and a couple of works have described various *Synechococcus* phylotypes [6,7]. Most recently, members of widespread clades such as SUP05 (*Ca*. Thioglobus spp), *Sulfurimonas* bacteria, and uncultivated SAR324 and Marinimicrobia have been studied from this and other dysoxic environments [8]. Here, we present a genome resolved metagenomic study of different depths in the Black Sea adding a total of 359 high-quality MAGs. The epipelagic and DCM strata show an overall marine-brackish community composition with predominance of microbes similar to the Mediterranean and the Caspian Seas [9,10]. The anaerobic compartment, that accounts for up to 80% of the total sea volume, had a much more exotic microbiota including various members of the microbial “dark matter”.

## Results and discussion

### Analysis of Metagenomic raw reads

We have generated metagenomic datasets from a Black Sea depth profile. Samples were collected along the Bulgarian coast at two stations (Fig. S1A). Sampling depth was guided by the physicochemical measurements (Fig. S1B) to cover representative temperature, oxygen, and chlorophyll-a values (Additional File 1). Thus, for St. 307, with the maximum depth of 1100 m, samples came from the near-surface at 5 m depth, the deep chlorophyll maximum (DCM) at 30 m, a sample from the redoxcline/pycnocline at 150 m and finally a sample at 750 m depth, corresponding to the euxinic water layer. Additionally, we collected a single near-surface sample (5 m) closer to the shore at station 301 with a maximum depth of 22.5 m. For each depth, we performed a first unassembled read analysis to obtain a rough taxonomic profile based on metagenomic 16S rRNA gene fragments against the SILVA database [11] (Fig. 1A) and the main predicted metabolic functions assessed by the SEED subsystems [12] (Fig. 1B).

**Fig 1.**
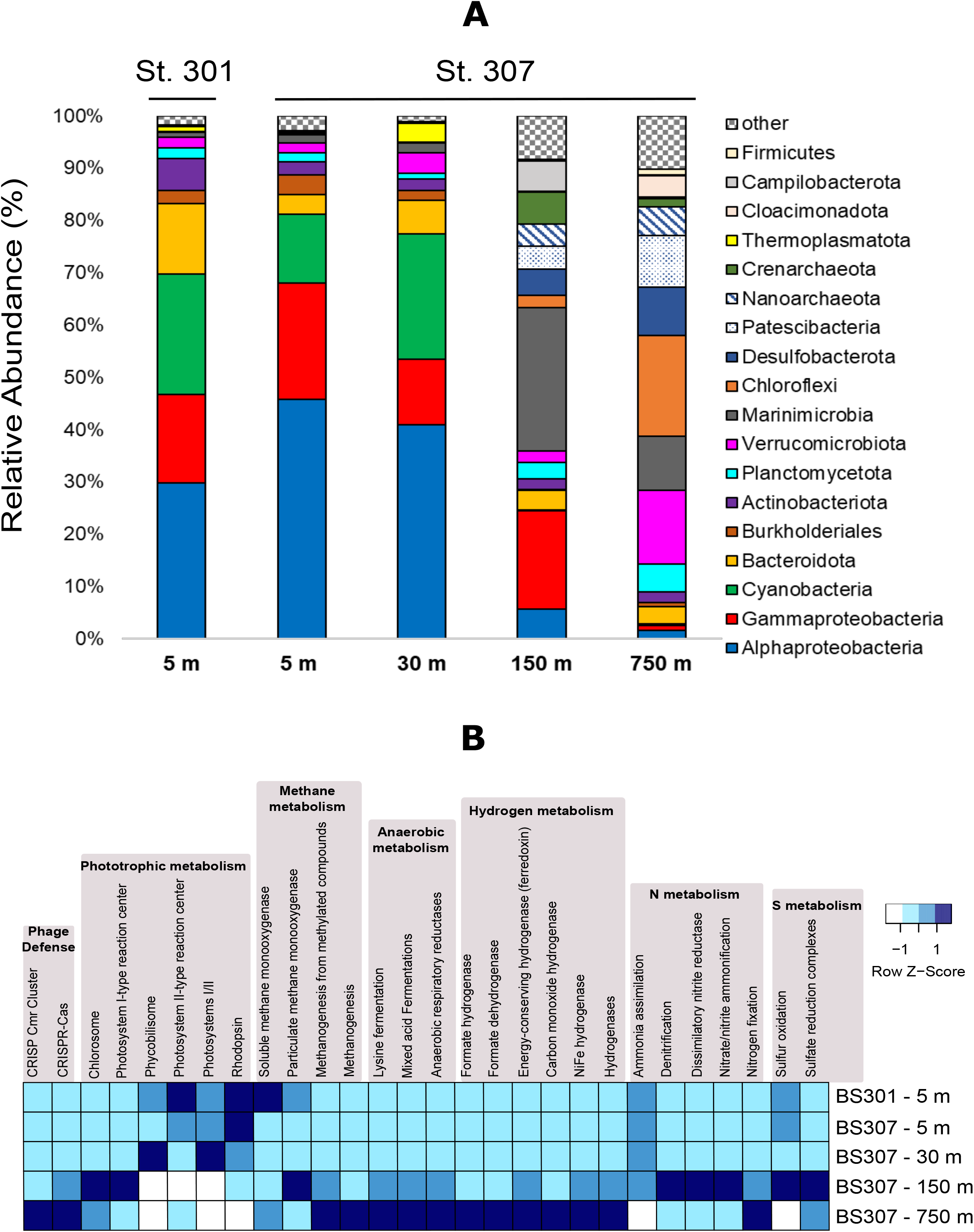
Black Sea phylum level 16S rRNA classification (A) and metabolic profiles assessed with SEED subsystems (B).

The oxic strata, surface and DCM, presented, at this rough level, very similar taxonomic composition (Fig. 1A). Alphaproteobacteria (orders SAR11, SAR116, Rhodobacterales and Rhodospirillales), Gammaproteobacteria (mostly orders SAR86 and Pseudomonadales), and picocyanobacteria (order Synechococcales) were the most abundant groups, representing > 70 % of total microbial biomass (assessed by total 16S rRNA classification). It must be highlighted the complete absence of the genus *Prochlorococcus* in all our Black Sea samples. The predominant subsystems of the oxic layer (Fig. 1B) were, as expected, associated with phototrophic lifestyles such as those from Synechococcales (photosystems/phycobilisomes) or photoheterotrophy with type-1 rhodopsin pumps (typical of SAR11, SAR86 or Flavobacteriales). In addition, ammonia was the preferred N source.

The taxonomic composition changed dramatically as we reached oxygen extinction in the pycnocline (150 m), where various taxa and microbial lifestyles coexisted, with a prevalence of anaerobic N and S related subsystems (Fig. 1B). Marinimicrobia (ca. 30 % of 16S rRNA assigned reads) and Gammaproteobacteria (ca. 20 %) were the dominant taxa of the redoxcline (Fig. 1A). Chemolithotrophs and anaerobes, such as SUP05 (*Ca*. Thioglobus spp.), Nitrosopumilaceae (aerobic archaeal ammonia oxidizers), Campylobacterota (dissimilatory nitrate reducer), Marinimicrobia (fermenters and hydrogen metabolizers), Nitrospirota (nitrite oxidizers) (N fixers), Chlorobi (anoxygenic photosynthesizers), Desulfobacterota (sulfate-reducers) and various associated streamlined microbes such as Patescibacteria and Nanoarchaeota appeared here.

The euxinic waters at 750 m showed an increase in fermentation, hydrogen metabolism, anaerobic respiratory reductases or methanogenesis pathways (Fig. 1B). Overall, we observed an increase in sulfate reducers (Desulfobacterota), Dehalococcoidia/Anaerolineae Chloroflexota and a huge diversity of accompanying microbiota providing hydrogen and fermentation by-products that conformed a syntrophic network fueling the sulfate reducers at the redox end. There were representatives from Omnitrophota and Kiritimatiellae (both classified inside Verrucomicrobiota according to SILVA standards [11], although Omnitrophota is classified as a single phylum according to GTDB [13]), Phycisphaerae Planctomycetota, Marinimicrobia, Nanoarchaeota, Patescibacteria, andCloacimonadota.

Finally, Halobacterota (Syntrophoarchaeia) and Crenarchaeota (Bathyarchaeia) minor representation (< 2 % of total microbial biomass assessed by 16S rRNA) showed that methanogenesis coexisted with sulfate reduction in these euxinic waters if in much more reduce fraction.

### MAGs recovered from the different samples

Automated binning followed by manual curation of generated bins allowed the recovery of 359 MAGs with > 50 % completeness and < 5 % contamination. Detailed stats of these MAGs are described in Table 1 (MAGs from oxic samples) and Table 2 (redoxcline/anoxic MAGs) and in individual MAG detail in Additional File 2. Genomes are also showed in an estimated genome size versus GC content plot in Fig. S2. The taxonomic nomenclature used in this work was based on GTDB (ref). To estimate the binning efficiency, we mapped the reads of each metagenome against the MAGs obtained for each sample at the thresholds of > 95 % of identity and > 50 bp of alignment lengths. The percentages of reads mapped to the MAGs varied between samples, being maximum in the redoxcline (50 %) and minimum in the euxinic sample (33 %). With regard to the oxic samples, the MAG recovery efficiency was ca. 50 % of the total reads mapped with the MAGs from the coastal epipelagic sample (BS301-5 m), 42 % for the off-shore epipelagic sample (BS307-5 m) and 34 % for the DCM sample (BS307-30 m). These recovery values are in the range of what was previously obtained for other aquatic environments [14].

**Table 1.** Summary statistics and features of Black Sea MAGs retrieved from 5 and 30 m samples.

| Phylum/Division | Taxonomic affiliation of MAGs (GTDB, Referenced groups) | n° of MAGs | Range Estimated Genome size (Mp) | Range GC (%) | Range median intergenic spacer (bp) | Av. Compl. (%) | Av. Cont. (%) |
| --- | --- | --- | --- | --- | --- | --- | --- |
| α-Proteobacteria | **g_Planktomarina (3)**, **g_Puniceispirillum (6)**, **g_Reyranelia (1)**, **f_Rhodobacteraceae (7)**, **o_Pelagibacterales (8)**, **o_Parvibaculales (5)**, **f_Puniceispirillaceae (6)**, **f_Nisaeaceae (1)**, **o_Rickettsiales (3)**, **o_Rhizobiales (2)**, **o_Rhodospirillales_A (1)**, **c_Alphaproteobacteria (3)** | 52 | 1-7.9 | 28-66 | 2-65 | 75.55 | 1.19 |
| γ-Proteobacteria | **g_Luminiphilus (11)**, **g_Litoricola (3)**, **g_Nevskia (1)**, **f_Methylophilaceae (5)**, **f_Porticoccaceae (1)**, **f_Pseudohongiellaceae (5)**, **f_Shewanellaceae (1)**, **o_Burkholderiales (2)**, **o_SAR86 (11)**, **o_Pseudomonadales (5)**, **c_Gammaproteobacteria (3)** | 51 | 1-4.1 | 31-69 | 1-86 | 70.34 | 1.29 |
| Bacteroidota | **f_Cryomorphaceae (4)**, **f_Flavobacteriaceae (23)**, **o_Flavobacteriales (8)**, **f_Balneolaceae (2)**, **f_Crocinitomicaceae (2)**, **c_Bacteroidia (1)** | 44 | 1.19-2.57 | 28-58 | 4-43 | 73.91 | 0.94 |
| Thermoplasmatota | **f_Poseidonaceae (6)**, **f_Thalassarchaeaceae (1)**, **g_Poseidonia (6)** | 13 | 1.82-2.35 | 37-58 | 25-38 | 80.67 | 0.30 |
| Actinobacteriota | **o_Nanopelagiales (1)**, **g_Aquiluna (1)**, **o_Actinomarinales (5)**, **f_Illumatobacteraceae (4)**, **c_Thermoleophilia (1)** | 13 | 1.23-2.22 | 32-71 | 2-23 | 76.79 | 1.45 |
| Cyanobacteria | **g_Synechococcus_C (10)** | 10 | 1.8-2.23 | 55-63 | 25-36 | 79.59 | 2.37 |
| Planctomycetota | **f_Planctomycetaceae (1)**, **o_Pirellulales; (4)**, **p_Planctomycetota (3)**, **g_Rubripirellula (1)** | 9 | 3-6.4 | 49-72 | 45-132 | 87.14 | 0.95 |
| Verrucomicrobiota | **o_Pedosphaerales (2)**, **o_Opitutales (1)**, **f_Puniceicoccaceae (4)**, **f_Akkermansiacae (1)** | 8 | 2-4.8 | 42-60 | 23-74 | 83.51 | 4.21 |
| Marinisomatota | **g_Marinisoma (3)** | 3 | 0.8-0.93 | 31-32 | 2-3 | 55.31 | 0.36 |
| Margulisbacteria | **c_ZB3 (1)** | 1 | 1.71 | 42.5 | 10 | 67.53 | 0 |

Parenthesis ( ) indicate the average value of each field in case of range values and number of MAGs in bold for each taxonomic affiliation. Taxonomic classification follows GTDB criteria. Marinisomatota includes former Marinimicrobia. Thermoplasmatota includes former Euryarchaeota group. d\_:Domain, p\_:Phylum, c\_:Class, o\_:Order, f\_:family, g\_:Genus, s\_: Species. Av. (Average), Compl. (Completeness), Cont. (Contamination).

**Table 2.**
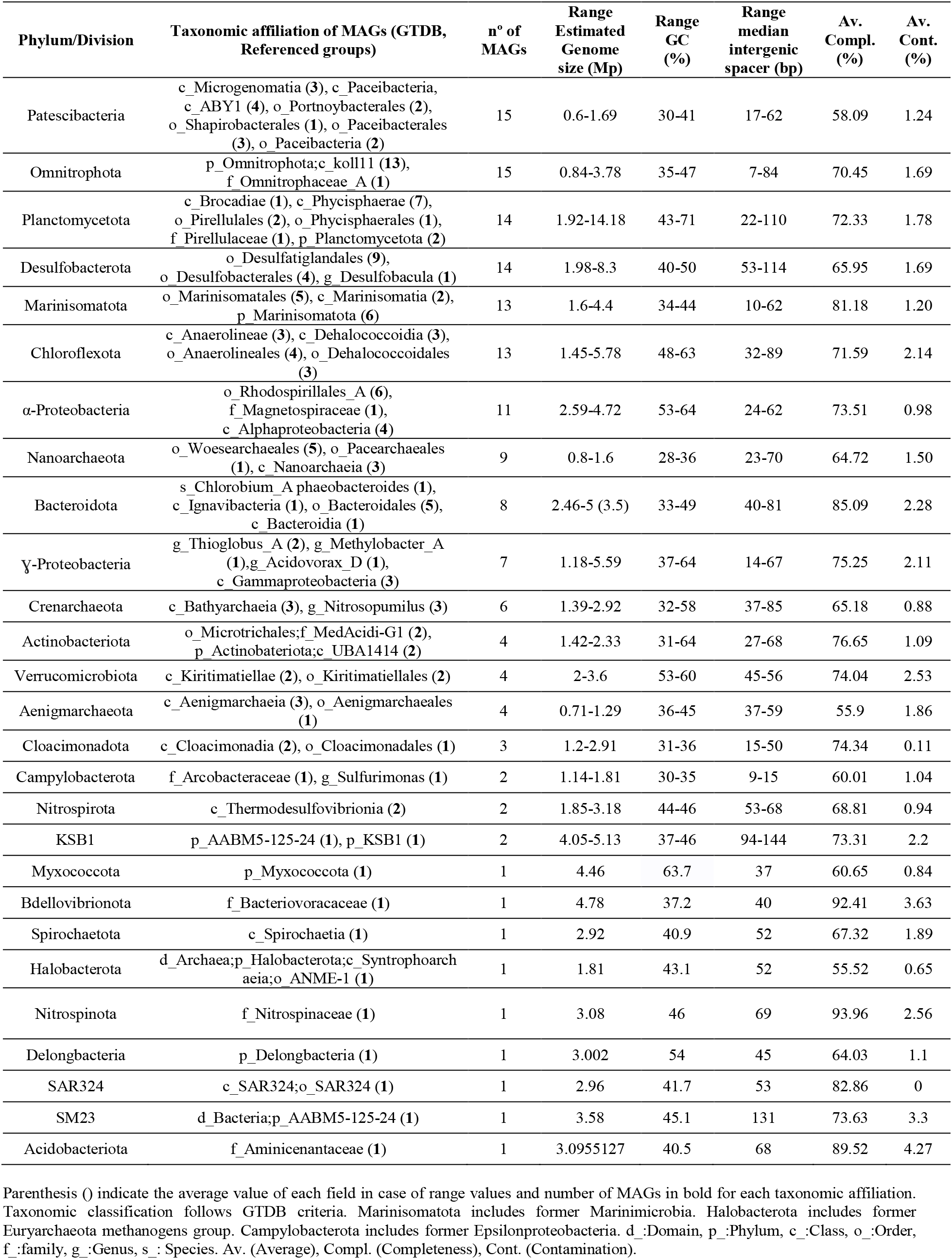
Summary statistics and features of Black Sea 150 and 750 m retrieved MAGs.

A Distance-based redundancy analysis (dbRDA) was conducted to statistically assess the main differences between different Black Sea strata (Fig. 2). To make such analysis we used the physicochemical measurements (Additional File 1), the metabolic abundance of each SEED subsystem (Fig. 1D) and the relative abundance of each microbial species retrieved as MAG and assessed with reads per Kb of Genome per Gb of metagenome (RPKGs), showed in Additional File 3.

**Fig. 2.**
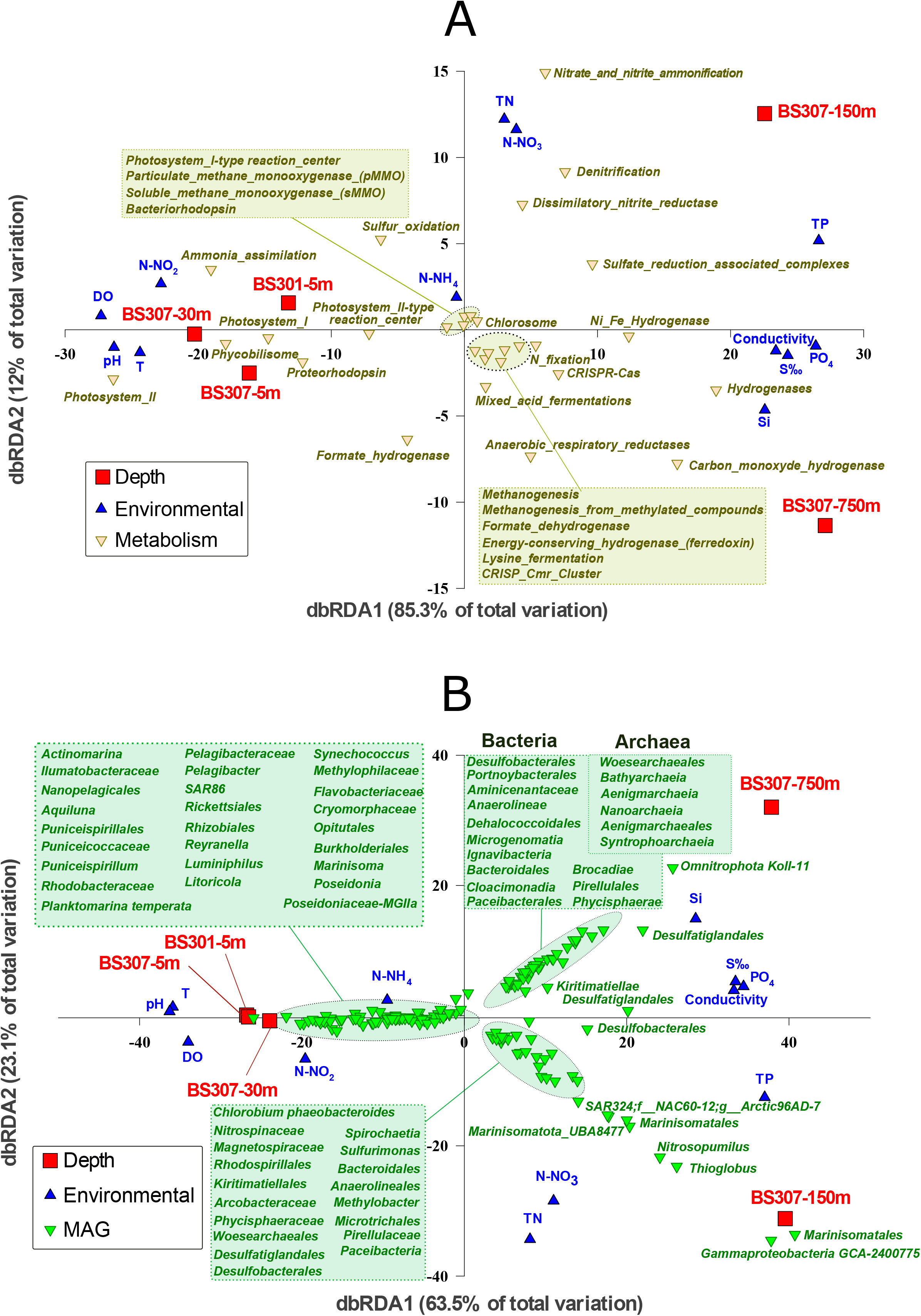
Black Sea dbRDA analysis between different samples (depths), environmental parameters and A) Metabolic pathways and B) MAG classification.

### MAGs from the epipelagic and DCM oxic strata

As expected, the statistical analysis conducted with the dbRDA (Fig. 2) grouped together the environmental variables of Temperature (T), dissolved oxygen (DO) or ammonia with photo(hetero)trophic lifestyles from well-known marine and brackish groups such as SAR11, SAR116 and Rhodospirillales (Alphaproteobacteria), SAR86 (Gammaproteobacteria), Thermoplasmatota (former marine group II Euryarchaeota), Synechococcales (*Synechococcus*) and Actinomarinales (Actinobacteria). As noted above, we must highlight a complete absence of *Prochlorococcus* spp., contrasted with a high abundance of various *Synechococcus* MAGs that affiliated with the marine clades I, III, IV, VI and WPC1 including isolates (KORDI-49, BL107, CC9902, WH 8016, WH 7805/7803, WH 8103/8102) [15]. The main Actinobacteria MAGs retrieved presented relatively small genome sizes (1.2-2.2 Mb), among which we must highlight the presence of 5 novel Actinomarinales (BS301-5m-G7, BS307-5m-G2, BS30m-G2/G3/G4) and a group of Ilumatobacteraceae genomes related to Caspian MAGs (Casp-actino5) [10]. The major SAR11 Alphaproteobacterial MAGs were eight novel Pelagibacterales that affiliated with the recently described groups Ia.1, IIaB/1and IIIa [16]. Remarkably, we obtained three novel MAGs from the order Rickettsiales. Another relevant Alphaproteobacteria clade from which we obtained MAGs was SAR116, with six MAGs affiliated to *Puniceispirillum* genus and five more were only classified as representatives of the family Puniceispirillaceae. A remarkable family that has shown a high abundance in Black Sea oxic waters is Flavobacteriaceae (23 MAGs), a group that was commonly detected in the Mediterranean [9] and the Baltic Seas [17]. In fact, various MAGs were related at GTDB genus level with MED-G11, MED-G14 MAGs and at the species level (ANI > 95 %) with MED-G20 Mediterranean Sea MAGs. Two MAGs also showed their closest relatives at the GTDB family level with Baltic Sea MAGs BACL11 and at the species level with BACL21. We also found five representatives from the clade OM43 (family Methylophilaceae) affiliating at the genus level to BACL14 Baltic Sea MAGs. Eleven MAGs belonged to the cosmopolitan Gammaproteobacteria SAR86, so far only classified at this order level. Other Gammaproteobacteria that co-occurred in these samples were MAGs with similarity to *Luminiphilus* (11 MAGs) and *Litoricola* (3 MAGs) genera. Another relevant taxon from marine systems was the former marine group-II Euryarchaeota (Thermoplasmatota according to GTDB taxonomy). We retrieved six genomes affiliating to the family Poseidoniaceae and other six to the genus Poseidonia. Only one genome was obtained affiliating to Thalassoarchaeaceae. Finally, three ultra-small (1 Mb of estimated genome size) Marinimicrobia MAGs were obtained from oxic metagenomes, which so far are classified by the GTDB as genus *Marinisoma*.

### Black Sea pycnocline MAGS

The redoxcline of the Black Sea presented the most metabolically diverse set of pathways among all analyzed samples (Fig. 2A). The main environmental variables that statistically grouped with the pycnocline were total nitrogen (TN) and nitrate, which were clearly associated with the different N cycle pathways that completed its biogeochemical cycle in this layer. The highest abundance of N pathways corresponded with denitrification (nitrogen gas as the final product), nitrate/nitrite ammonification and dissimilatory nitrate reduction (with ammonium as the final product), but the N cycle was also completed with ammonia oxidation and N fixation pathways detected both in total reads and MAGs (see below). Nonetheless, various other metabolisms coexisted in this thin layer where oxygen is extinguished. We noticed the presence of anoxygenic photosynthesis, exemplified by MAG BS150m-G13 showing > 99 % of ANI with *Chlorobium phaeobacteroides*, a green sulfur bacterium (GSB) originally isolated from the Black Sea [18] (GCA_000020545.1), that was undergoing a nearly monoclonal bloom (Fig. S3).

Chemoautotrophy was observed in *Thioglobus* sp. BS150m-G29 and G33 MAGs, both of which are novel representatives of the SUP05 clade which performs a wide variety of metabolisms including S oxidation and C fixation and with only 80 % of ANI with its closest relative (*Ca*. Thioglobus autotrophicus EF1) [19]. It appears that this is a case of a single species (recruiting at > 95 % of nucleotide identity) abundant (> 70 RPKG, Fig. S4) in the Black Sea redoxcline. Methane oxidation (*Methylobacter* sp. BS150m-G31) and ammonia oxidation were also key metabolisms observed in this layer (*Nitrosopumilus* spp. BS150m-G38/39/40). Nitrite oxidation was detected in Nitrospinaceae BS150m-G45.

Denitrification was frequently detected among pycnocline MAGs, although complete denitrification including the last step involving conversion of nitrous oxide into nitrogen gas (*nosZ* gene) was seen only in five MAGs (Marinimicrobia BS150m-G46/G47/G71, unclassified Alphaproteobacteria BS150m-G7/G9, Rhodospirillales BS150m-G4/G10, *Sulfurimonas* sp. BS150m-G26 and unclassified Gammaproteobacteria BS150m-G28/30/32). Dissimilatory nitrate reduction to ammonium (*nrfAH* genes) was far more restricted and found in Marinimicrobia BS150m-G46, Campylobacterota (*Sulfurimonas* sp. BS150m-G26) or Bacteroidales BS150m-G15. Nitrate reduction through *nirB* gene was much more widely detected including in all Alphaproteobacteria MAGs and various Gammaproteobacteria members. N fixation (nifDK dinitrogenase subunits) was detected in only two MAGs (*Chlorobium phaeobacteroides* and Nitrospirota, BS150m-G55/G56 respectively).

Dissimilatory sulfate reduction and oxidation (*dsrAB* genes) showed up already in this sample in various genomes such as Desulfobacterota, Nitrospirota MAGs, Planctomycetota (Pirellulaceae BS150m-G36) (already mentioned above), Chloroflexota (Anaerolineales BS150m-G18), Alphaproteobacteria (Rhodospirillales BS150m-G3/G4/10/G11), *Chlorobium phaeobacteroides* MAG and Gammaproteobacteria (*Ca*. Thioglobus and Gammaproteobacteria BS150m-G28 MAGs). It must be noted that, among the main features of this habitat, there was the simultaneous activity of sulfate-reducing and sulfide-oxidizing microbes forming part of the same ecological niche, a process known as cryptic sulfur cycle [20]. However, low O_2_ concentrations (0.87 mg/L) and low ratio (0.16) of peroxidase/recA genes (1.5 in oxic datasets) clearly demonstrate the microaerophilic/anoxic nature of this habitat.

We also compared our pycnocline dataset with previously available metagenomes from the redoxcline from Cariaco Basin (Venezuela) [21] (Fig. S5). Overall, it seems that Marinisomatota/Marinimicrobia and Gammaproteobacteria chemolithotrophic groups are the most abundant key players of these two marine redoxclines, accounting for more than 50 % of total microbial biomass (Fig. S5A). However, it must be noted that only a few species retrieved as MAGs from the Black Sea were detected in such a similar habitat (Fig. S5C). Among them, two chemolithotrophic Gammaproteobacterial representatives (*Ca*. Thioglobus and a novel species BS150m-G30 classified only at the order level as o GCA-2400775 by GTDB), sulfate reducers (Desulfatiglandales), denitrifying and hydrogen-producing Marinimicrobia (three species) and one Actinobacteria (a novel species from the marine MedAcidi-G1 group). Apart from their metabolic potential fitting with microbial lifestyles from pycnocline layers, these species could play key roles in other marine redoxclines and oxygen minimum zones (OMZs), as their detection in two largely separated biomes with different salinities (ca. 2 % in the Black Sea and 3.5 % in the Cariaco Basin) indicate a widespread distribution in oxygen-depleted marine niches.

### Euxinic Black Sea MAGs and the “microbial dark matter”

The main environmental variables grouping with the mesopelagic sample of 750 m were PO_4_ and Si, both of which are solubilized in anoxic layers and diffuse from the sediment layer. Salinity also increased up to 2.2 % in these euxinic waters. There is the expected predominance of sulfate reduction pathways, as carried out by Desulfobacterota MAG representatives (Desulfatiglandales BS750m-G47-G51 and BS750m-G54/G56, Desulfobacterales BS750m-G52/G53/G55), which perform the dissimilatory sulfate reduction pathway (*dsr* genes).

Methanogenesis was very diluted in these waters but still detectable, albeit we found the complete pathway in MAG *Ca.* Syntrophoarchaeum BS750m-G82 and most of the genes except for the key enzyme, Methyl-coenzyme M reductase (*mcr*) in Bathyarchaeota BS750m-G27/G28 MAGs as well. The latter showed various mixed-acid fermentation pathways including the formation of H_2_ and CO_2_ (via the formate hydrogenlyase), formate (pyruvate-formate lyase), alcohol (alcohol dehydrogenase) or lactate (lactate dehydrogenase). It must be noted the potential capability of performing reverse methanogenesis, or anaerobic methane oxidation (ANME) by the abovementioned archaeon (MAG *Ca*. Syntrophoarchaeum BS750-G82). Heterodisulfide reductase genes (*HdrABC*), which are involved in the last step of methanogenesis by reducing CoB-CoM heterodisulfide, were detected in Bathyarchaeota and *Syntrophoarchaeum* MAGs. However, these genes were also found in Cloacimonadota, candidate division KSB1, *Ca*. Aminicenantes, Omnitrophica, Desulfobacterota, Planctomycetes, andChloroflexi MAGs as well as in the unassembled reads (being completely absent from oxic datasets), suggesting that these electron transfer complexes are not exclusive of methanogens. As seen by the dbRDA, we also noted a global predominance of mixed-acid fermentation pathways (with ethanol, lactate, acetate, formate or CO_2_/H_2_ as products) and hydrogen uptake hydrogenases that couple with sulfate, fumarate, CO_2_ or nitrate reduction, thus conforming a complex syntrophic network of microbes. This networking of syntrophic microbes (considered here as interspecies H transfer) includes the abovementioned uncultured taxa plus accompanying streamlined members of the “microbial dark matter” such as Omnitrophota, Patescibacteria (*Ca*. Microgenomates, Portnoybacteria, Paceibacteria) or Nanoarchaeota (*Ca*. Aenigmarchaeota, Woesearchaeota, Pacearchaeota), groups from which we also obtained MAGs (see Table 2). Various types of hydrogenases and hydrogen metabolism pathways grouped with the 750 m mesopelagic sample in the dbRDA plot (Fig. 2) and were found in the vast majority of microbes inhabiting this sulfide enriched waters, including NAD-reducing bidirectional (*hox* genes) and uptake hydrogenases (*hup* genes), NiFe (*hyp* genes) and FeFe (*hym* genes) hydrogenases, Coenzyme F420-reducing hydrogenases or carbon monoxide induced hydrogenases (*CooHL* genes), all of which showed the highest gene/ recA ratios (from 0.2 in *hym* genes to 1-2 for *hyp* and *hoxF*) in euxinic waters.

It was remarkable the presence of two Actinobacteria MAGs (BS750m-G1/G2) in these sulfide-rich waters. These yet unclassified members have their highest resemblance with MAGs retrieved from groundwater aquifers (Actinobacteria bacterium CG08_land_8_20_14_0_20_35_9, classified as UBA1414 by GTDB) and have very small GC content (31-34 %) and predicted genome sizes (ca. 1.4-1.6 Mb). Their genomes presented various mixed-acid fermentative pathways associated with the production of ethanol (alcohol dehydrogenase), lactate (lactate dehydrogenase), formate (pyruvate-formate lyase) and H/CO_2_ (formate hydrogen lyase). They also showed an active hydrogen metabolism with various NiFe hydrogenases including Coenzyme F420-reducing hydrogenase, *hyp* genes and HyaA COG1740 355 Ni-Fe-hydrogenase I. Another remarkable group of microbes was Omnitrophota, from which we obtained 15 MAGs with variable estimated genomes sizes (from 1 to 3 Mb). For instance, the most abundant MAG retrieved from our samples (BS750m-G77) presented a small predicted genome size (ca. 1.2 Mb) and was an obligate fermenter (mainly producing ethanol, H_2_/CO_2_ and lactate). Another group of streamlined members of the microbial dark matter were Aenigmarchaeota (BS750m-G24/36/81/83/) and Nanoarchaeota (BS750m-G11/13/70) MAGs, which had estimated genome sizes of 1-1.5 Mb. Among their metabolic potential, they were also mixed-acid fermenters, including lactate or H_2_/CO_2_ as fermentation by-products, which would fuel the sulfate reducers, conforming a syntrophic network with the rest of mixed-acid fermenters. Finally, another set of microbes of small genome sizes (0.6-1.6 Mb) were Patescibacteria (former Candidate Phyla Radiation). We must highlight the presence of the protein VirB4, associated with type IV secretion systems that work as injectors into host cells [22], in Ca. Microgenomates BS750m-G73/74, Ca. Paceibacteria BS750m-G71/75 and Ca. Portnoybacteria bacterium BS750m-G76. These proteins were unique for these microbes in the entire euxinic waters, which suggests a parasitic lifestyle from which these Patescibacteria could translocate nutrients, proteins, and DNA from or to a putative host [22].

### Similarities between Black Sea datasets assessed by read and recruitment analysis

To assess the representativity of our samples we also compared the reads between our Black Sea datasets and those from a former sampling campaign [5,8], deposited into the NCBI under bioproject PRJNA649215, observing a clear 16S rRNA taxonomy and read clusterization between samples (Fig. S6). Among all of the MAGs retrieved from this work, we selected the 30 most abundant MAGs (> 10 RPKGs in any of the recruited samples) from the oxic, redoxcline and anoxic waters and recruited them at > 95 % of identity (species level) on all metagenomes (Fig. 3). The rest of our MAGs abundance among all datasets is shown in detail in Additional File 3. As expected, emblematic key players of the oxic waters harboring a phototrophic/photoheterotrophic lifestyle such as *Ca*. Pelagibacter, *Ca.* Actinomarina, SAR86, *Synechococcus* or Flavobacteriaceae were detected at high numbers in all oxic metagenomic datasets. Next, we also showed the main ecological drivers of the redoxcline layer, which included chemolithotrophic S oxidizers and C fixers such as *Ca.* Thioglobus and dissimilatory nitrate reducers such as *Sulfurimonas*, both of which were recently analyzed members by previous publication [8]. The redoxcline also showed some other ecologically relevant nitrate reducers such as Bacteroidales MAGs, ammonia oxidizers such as *Nitrosopumilus* spp, sulfate reducers Desulfatiglandales and Desulfococcales novel species and several Marinimicrobia representatives which are specialized in the H metabolism, denitrification and mixed-acid fermentation. Finally, another set of MAGs were detected among all euxinic strata. These included various other sulfate-reducers, their associated microbiota performing mixed-acid fermentations and H metabolism in syntrophism (Cloacimonadetes, Woesearchaeales, *Ca*. Aminicenantes) and the only MAG able to perform methanogenesis and ANME, a *Syntrophoarchaeum* that showed a remarkable abundance in 1000 and 2000 m metagenomic datasets, suggesting that methane metabolisms indeed coexist with the sulfate reduction and all associated microbial fermenters and H_2_ scavengers in a complex syntrophic network.

**Fig. 3.**
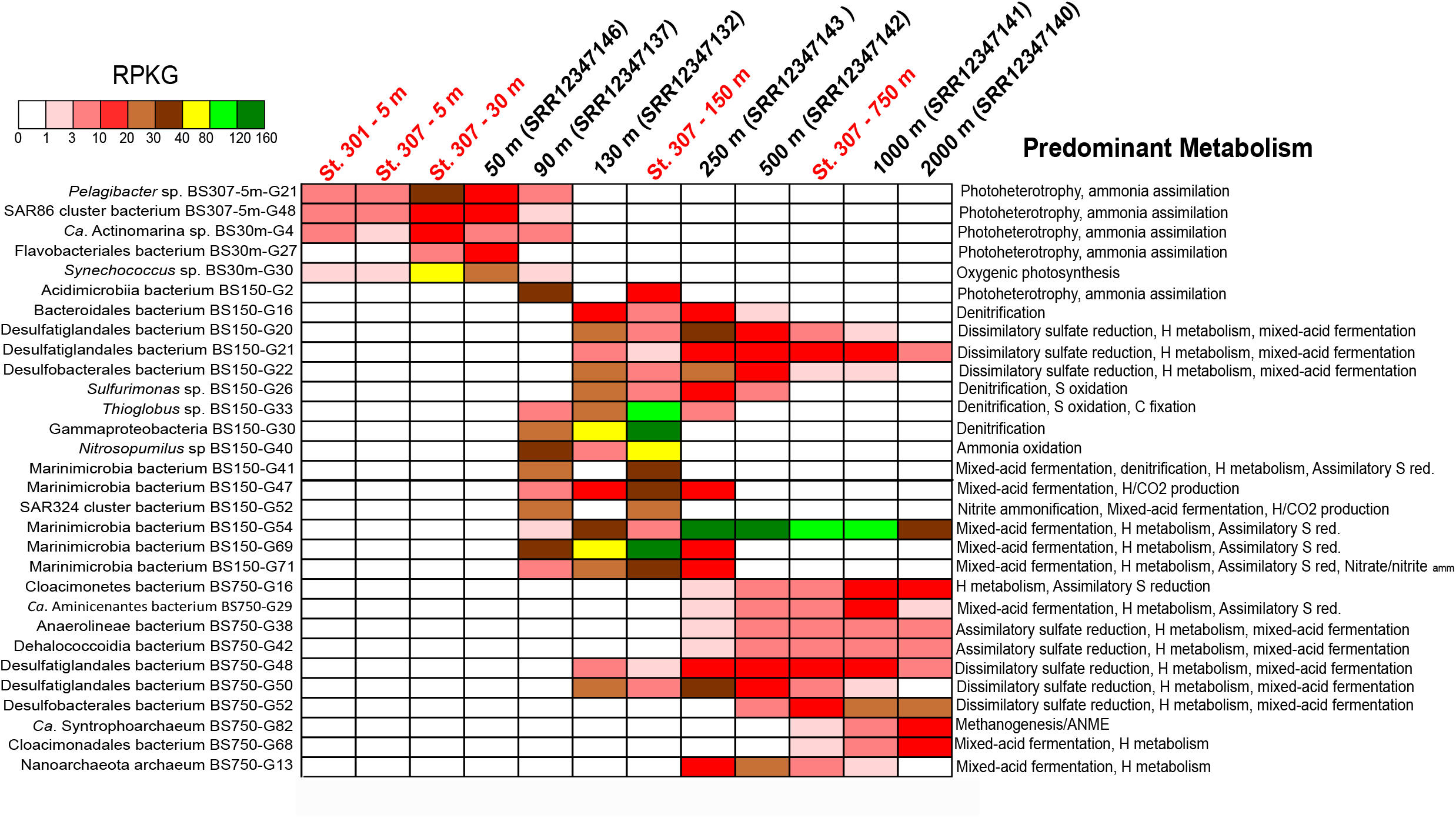
Recruitment analysis of the 30 most abundant Black Sea MAGs retrieved from our datasets (in red) and detected at highest values at various Black Sea metagenomes from the NCBI (Bioproject PRJNA649215). Reads were recruited at > 95 % of identity and > 50 bp of alignment lengths. The predominant metabolism is characteristic of each MAG and was detected in the genome and assessed by the literature.

## Conclusions

The present study, together with others from the same Black Sea [5,8] (bioproject PRJNA649215) and those recent ones from the Cariaco Basin in Venezuela [21,23] show a first glimpse on the microbiome of anoxic marine water columns. The redoxclines of these habitats show a convergence of various metabolisms at a time, among which we encounter anoxygenic photosynthesis such as that one observed in *Chlorobium phaeobacteroides* [24], ammonia oxidation by *Nitrosopumilus* spp., chemolithotrophic metabolisms carried by Gammaproteobacteria such *Ca*. Thioglobus, one of the most abundant and versatile players transitioning between oxic-anoxic regimes [19,25]. This microbe has been detected both in Cariaco and Black Sea basins [8] and its adaptive metabolism, which includes physiological adaptations to the oxic-anoxic growth [25] and its wide set of metabolic tools has led it to colonize these zones with a large contribution to S and N biogeochemical cycles as a denitrifier, sulfur-oxidizing and C fixer chemolithotroph. The simultaneous activity of sulfate-reducing and sulfide-oxidizing microbes in these habitats has led to a term known as the “cryptic sulfur cycle” [20]. One of the most abundant microbes detected from these zones is Marinimicrobia, which are specialists of both oxic and anoxic waters [26,27], able to perform denitrification and various fermentations and H_2_ metabolism and were recently labeled as organoheterotrophs with specific molybdoenzymes to preserve energy from sulfur cycle intermediates [8].

As we approach the euxinic mesopelagic and bathypelagic waters, we tend to encounter a fundamental domination of sulfate reduction coupled with a complex variety of syntrophic networks that feed the ecosystem. In this sense, Desulfobacterota is a good example of a syntrophic phylum that could be able to accept electrons from other electron donors, as noted previously in marine sediments [28]. It appears that the extremely high abundance of sulfate-reducers in the Black Sea has displaced methanogens, which are present in the water column but at low numbers, having obtained Bathyarchaeota [29,30] representatives and a single example of a *Syntrophoarchaeum* from this study. In fact, this last microbe could be performing reverse methanogenesis or anaerobic methane oxidation (ANME) in the system. Recently, members of this newly identified species have shown the complete oxidation of butane during the anaerobic methane oxidation process [31]. However, it appears that the competition between methanogens and sulfate-reducers for acetate is dominated by the latter, which also take a fundamental role in the syntrophic network by uptaking all the H_2_ produced by the fermenters. Among all of the associated microbiota, we must highlight the presence of Cloacimonadota phyla (previously known as WWE1), which have shown up in meromictic lakes as important carbon and sulfur recyclers [32] and appear to degrade propionic acid in syntrophic networks in bioreactors [33]. Novel microbes from lineages such as Cloacimonadota (WWE1), Marinisomatota (SAR406), Omnitrophicaeota (OP3), Bacteroidales, Kiritimatiellae, Anaerolinea/Dehalococcoidia formed a very complex syntrophic network where mainly mixed-acid fermentations with lactate, ethanol, formate, succinate, hydrogen and CO_2_ were formed as final products. Other exotic members of this system included the uncultured microbial dark matter, such as Patescibacteria and Nanoarchaeota [34–37]. In particular, we have stumbled upon various members of Aenigmarchaeota [38] and Woesearchaeales that showed streamlined genomes.

## Methods

### Sampling, DNA extraction, physical and chemical profiles measurement

Samples for metagenome analyses were collected from St. 301 (5 m deep) and St. 307 (5, 30, 150, and 750 m deep) in October 2019 (coordinates 43.155517 N 28.005383 E and 43.1696 N and 29.001283 E, respectively). Up to 6.9 L of seawater from each sampling depth were filtered through a series of 20 μm Nylon Net filters (Millipore), 5 μm polycarbonate membrane filters (Millipore), and 0.22 μm SterivexTM Filter Units (Merck). DNA was then extracted using standard phenol-chloroform protocol [39]. In short, Sterivex filters were treated with CTAB lysis buffer and then treated with 1 mg ml^−1^ lysozyme and 0.2 mg ml^−1^ proteinase K (final concentrations). Then nucleic acids were purified with phenol/chloroform/isoamyl alcohol and chloroform/isoamyl alcohol.

### Sequencing, assembly and read annotation

The five samples were sequenced in one lane of Illumina HiSeq X Ten PE 2X150 bp (Novogene company), which provided ca. 180-200 million clean reads and 24 Gb of output for each sample. Samples were individually assembled using IDBA-UD [40] with the parameters --pre_correction, --mink 50, --maxk 140, --step 10.Sub-assemblies of 20 million reads were done in each sample to retrieve some of the most abundant microbes that assembled poorly i.e. we reduced the total number of reads to obtain more complete bins of these representatives (e.g. Pelagibacterales, Actinomarinales, *Synechococcus*, Marinimicrobia or *Thioglobus* spp.). Annotation of contigs was assessed using Prodigal [41] for ORF prediction and then BLAST (nr database) using Diamond for functional annotation [42]. Proteins were annotated with latest nr, COG [43], and TIGFRAM [44] to provide the most updated taxonomy. Features like tRNAs and rRNAs were detected with tRNAscan [45] and ssu-align [46], respectively.

### Binning, classification, and MAG statistics

Binning procedure was performed as follows: a first manual inspection was done assigning a hit (based on BLAST against Nr) to each CDS, which allowed us to classify contigs taxonomically into different phyla. Then, an initial binning step was applied for each set of contigs assigned to each phylum with METABAT2 [47] using coverage in the different samples. Afterwards, further manual inspection of contigs was applied using GC content, coverage and tetranucleotide frequencies to refine the bins [48,49]. Finally we only used MAGs with < 5 % contamination and > 50 % of completeness based on CheckM estimations [50]. MAGs were taxonomically classified according to the latest version of GTDB-tk and the database release89 [13] and whenever we could we used class, order, family, genus or species names for all of them.

### 16S rRNA read classification, hierarchical cluster read analysis, read functionality

The 16S rRNA gene reads were detected in a subset of 20 million reads from each metagenome. We first obtained candidates using USEARCH [51] with RefSeq 16S rRNA as database and then these putative 16S rRNA were confirmed using ssu-align[46]. Then, a BLASTN was performed against the SILVA database [11] (SILVA_138_SSURef_Nr99_Tax_silva from December 2019) to provide a taxonomic classification. Hierarchical cluster analysis (dendrograms) of different metagenomic samples with k-mer=21 bp was assessed with SIMKA [52] and Bray-Curtis indexes of presence/absence were obtained. Subsets of 20 million reads of each metagenomes were analyzed with BLASTX against SEED [12] database using Diamond [42], with parameters more-sensitive, max-target-seqs 1, e-value 0.00001 > 50 bp of alignment length and > 50 % identity. The top hits were analyzed in search of specific genes and pathways based on the SEED database. Hits were normalized by the total number of hit counts for each sample and a row Z-score was calculated to assess statistical differences between samples for each metabolic pathway.

### Relative abundance of MAGs

To estimate the relative abundance of the recovered genomes in various datasets we performed read recruitment and mapping, which was assessed considering BLASTN hits of the metagenomic reads against each MAG at > 95 % identity and 50 bp of alignment length thresholds, as indication of belonging to the same species. A microbe was considered present in a metagenome if it was detected at > 1 RPKG (Reads per Kb of Genome per Gb of Metagenome). All relative abundances of our MAGs on Black Sea datasets are shown in Additional File 3. Datasets used for recruitment included Black Sea (PRJNA649215) and Cariaco Basin (PRJNA326482).

### Redundancy analysis (RDA) of environmental variables, MAGs, and metabolic processes

Distance-based redundancy analysis (dbRDA) analysis was performed to describe the ordinations of the main MAGs and metabolic processes in an environmentally constrained space [53] and conducted with the R package vegan [54]. Environmental matrixes were constructed with 12 environmental variables for the 5 Black Sea samples. Each matrix was square-root transformed and normalized and subsequently transformed to Euclidean resemblance matrix. On the other hand, we constructed the other two matrixes with the recovered MAGs and metabolic processes with 30 selected metabolic processes. Both were standardized and square root transformed before performing a Bray–Curtis dissimilarity resemblance matrix. Two dbRDA were obtained, the first one comprising the recovered genomes (MAGs) matrix using environmental matrix as predictor variable, and a second one based on the metabolic processes matrix using environmental matrix as predictor variable.

### Availability of data and materials

All metagenomes and reconstructed genomes derived from this work are publicly available under the NCBI Bioproject PRJNA638805. Metagenomes were deposited to NCBI-SRA with the accession numbers SRR12042682-SRR12042686.

## Declarations

## Acknowledgements

This work was supported by grants “VIREVO” CGL2016-76273-P [MCI/AEI/FEDER, EU] (cofounded with FEDER funds) from the Spanish Ministerio de Ciencia e Innovación and “HIDRAS3” PROMETEO/2019/009 from Generalitat Valenciana. FRV was also a beneficiary of the 5top100-program of the Ministry for Science and Education of Russia. PJC-Y was supported by APOSTD/2019/009 Post-Doctoral Fellowship from Generalitat Valenciana. This research has been carried out thanks to the International Bilateral Project between the Italian National Research Council and the Bulgarian Academy of Science (CNR-BAS), and in the framework of the National Science Program “Environmental Protection and Reduction of Risks of Adverse Events and Natural Disasters,” approved by the Resolution of the Council of Ministers No. 577/17.08.2018 and supported by the Ministry of Education and Science (MES) of Bulgaria (Agreement No. D01-230/06.12.2018).

## Author contributions

FR-V, CC, PJC-Y and SM conceived the study. ND, VS, NS and SM performed metadata analysis and sample collection. PJC-Y and JR-G performed DNA extraction. PJC-Y, AP, MM, JH-M analyzed the metagenomic data. PJC-Y, FR-V and MM wrote the manuscript. All authors read and approved the manuscript.

## Consent for publication

All authors have read and commented the manuscript and consent the publication.

## Availability of data and material

All data derived from this work is publicly available in the NCBI-Genbank databases.

## Ethical approval and consent to participate

This article does not contain any studies with human participants or animals performed by any of the authors.

## Supplementary Figure legends

**Fig S1.**
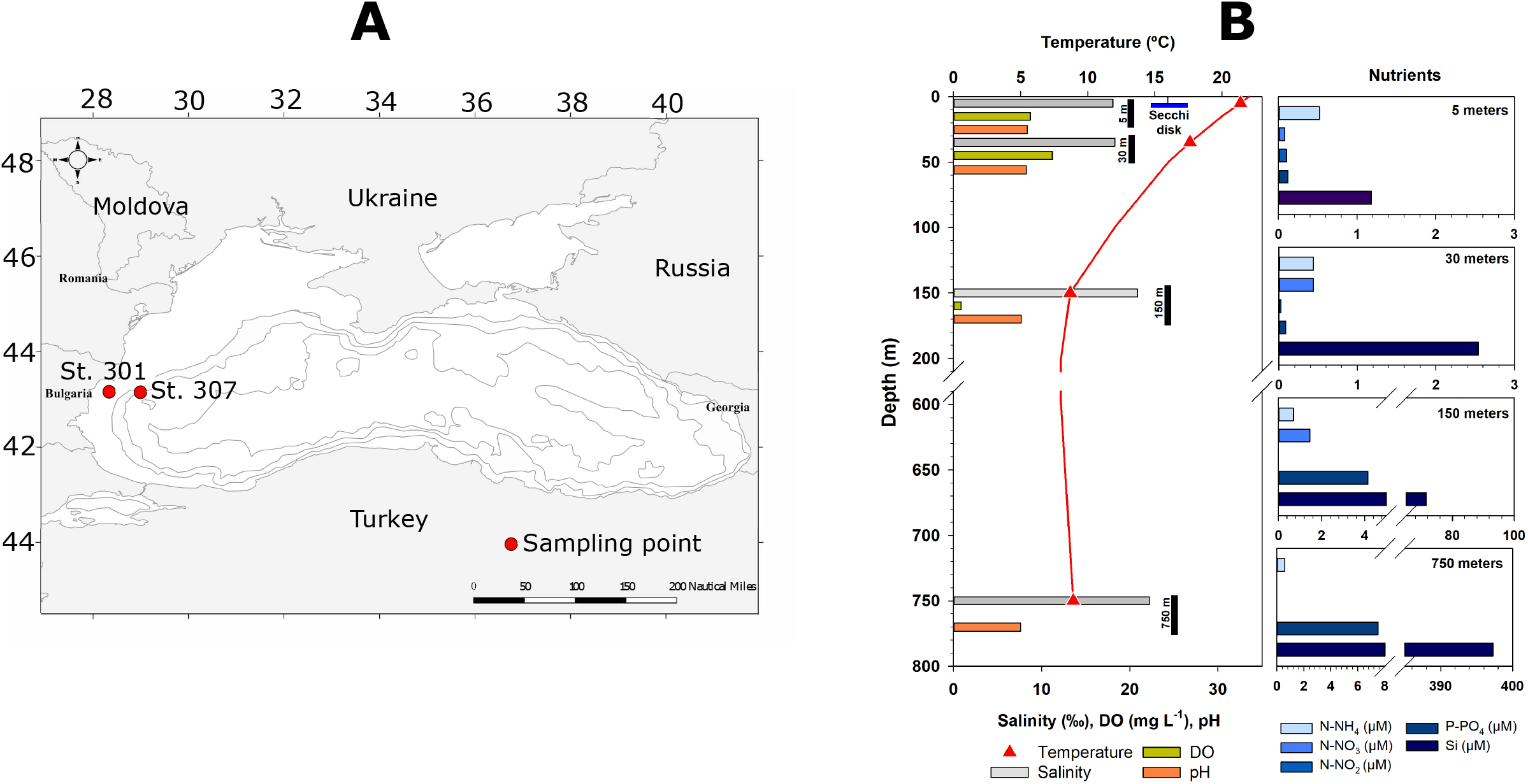
Black Sea sampling points (A) and physicochemical profiles (B). Each environmental measurement is color-coded.

**Fig S2.**
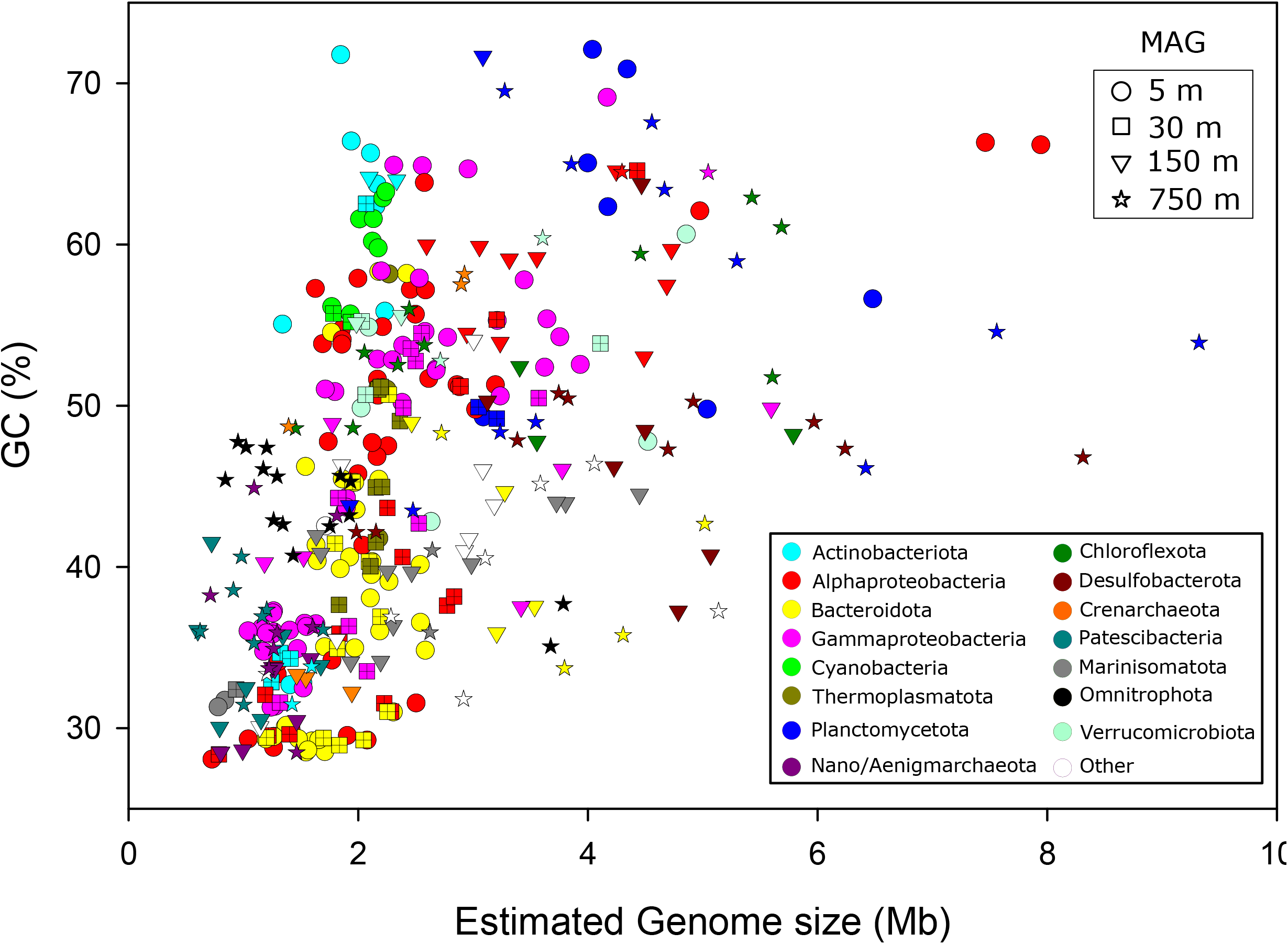
Estimated genome size (Mb) versus GC content of all Black Sea MAGs retrieved in this work. Shape indicates the depth at which the MAG was recovered. MAGs are color-coded at the phylum level.

**Fig. S3.**
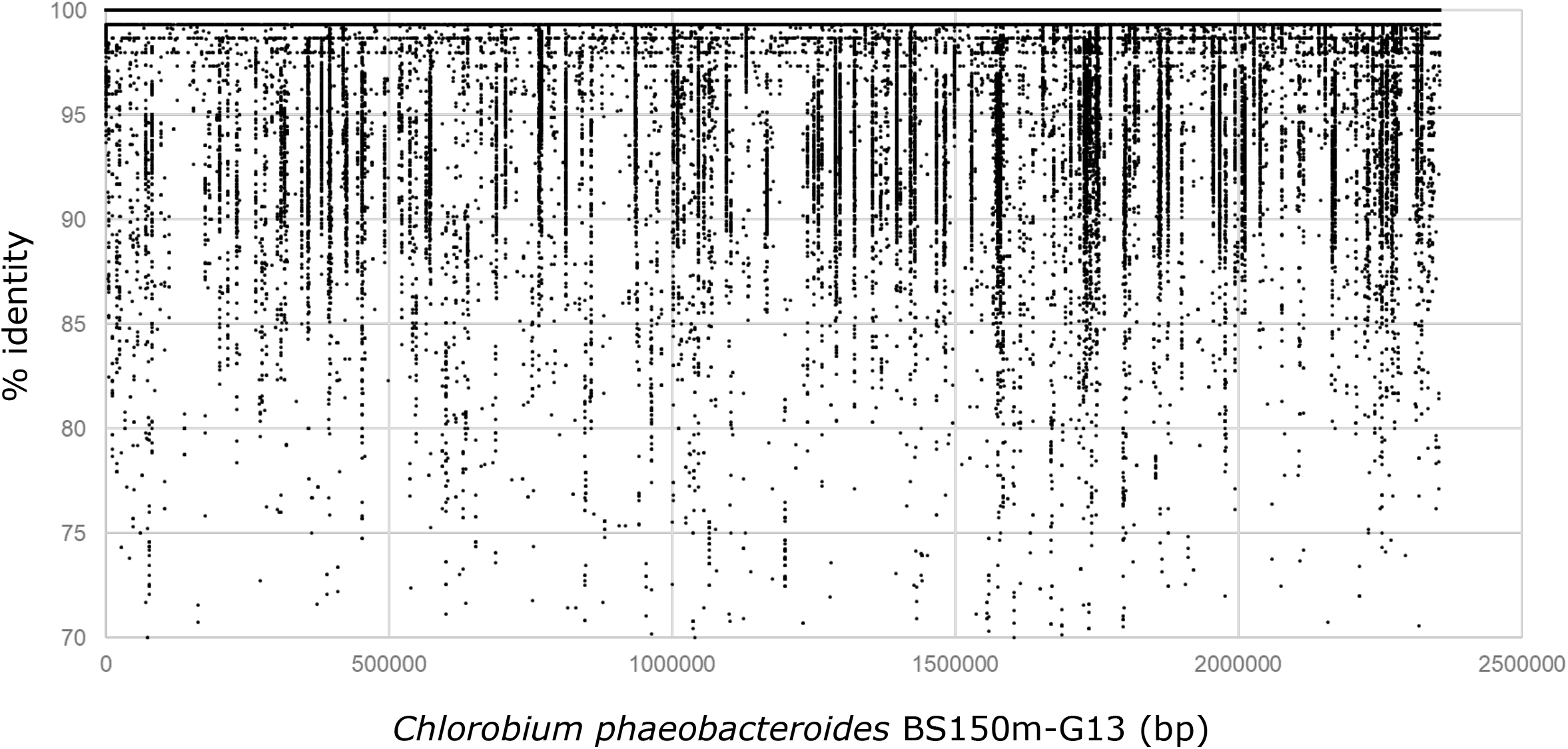
Recruitment plot of *Chlorobium phaeobacteroides* BS150m-G13 from the Black Sea 150 m pycnocline metagenome. Each dot represents a mapped read at > 95 % of identity and > 50 bp of alignment lengths.

**Fig. S4.**
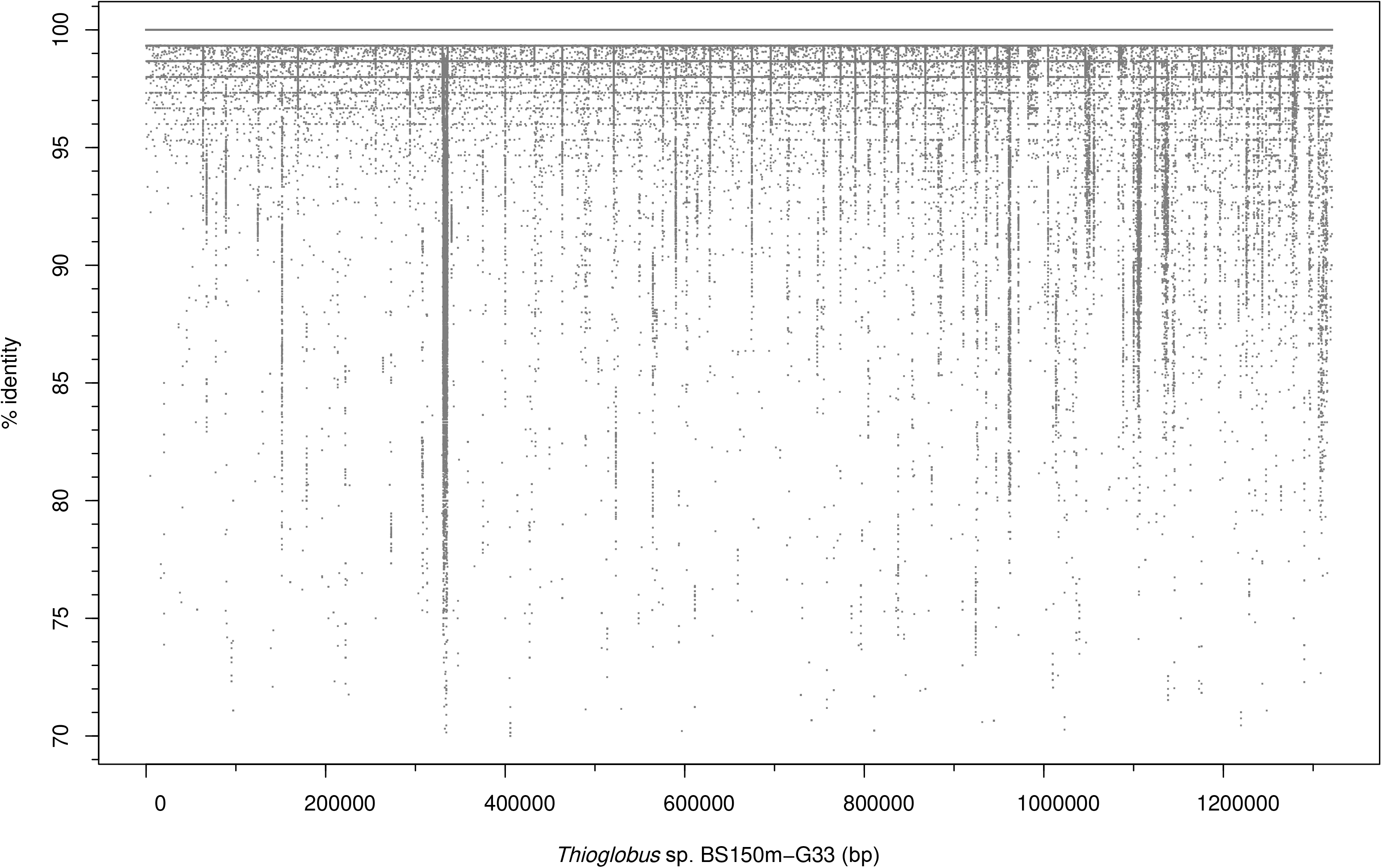
Recruitment plot of *Thioglobus* sp. BS150m-G33/G29 from the Black Sea 150 m pycnocline metagenome. Each dot represents a mapped read at > 95 % of identity and > 50 bp of alignment lengths.

**Fig. S5.**
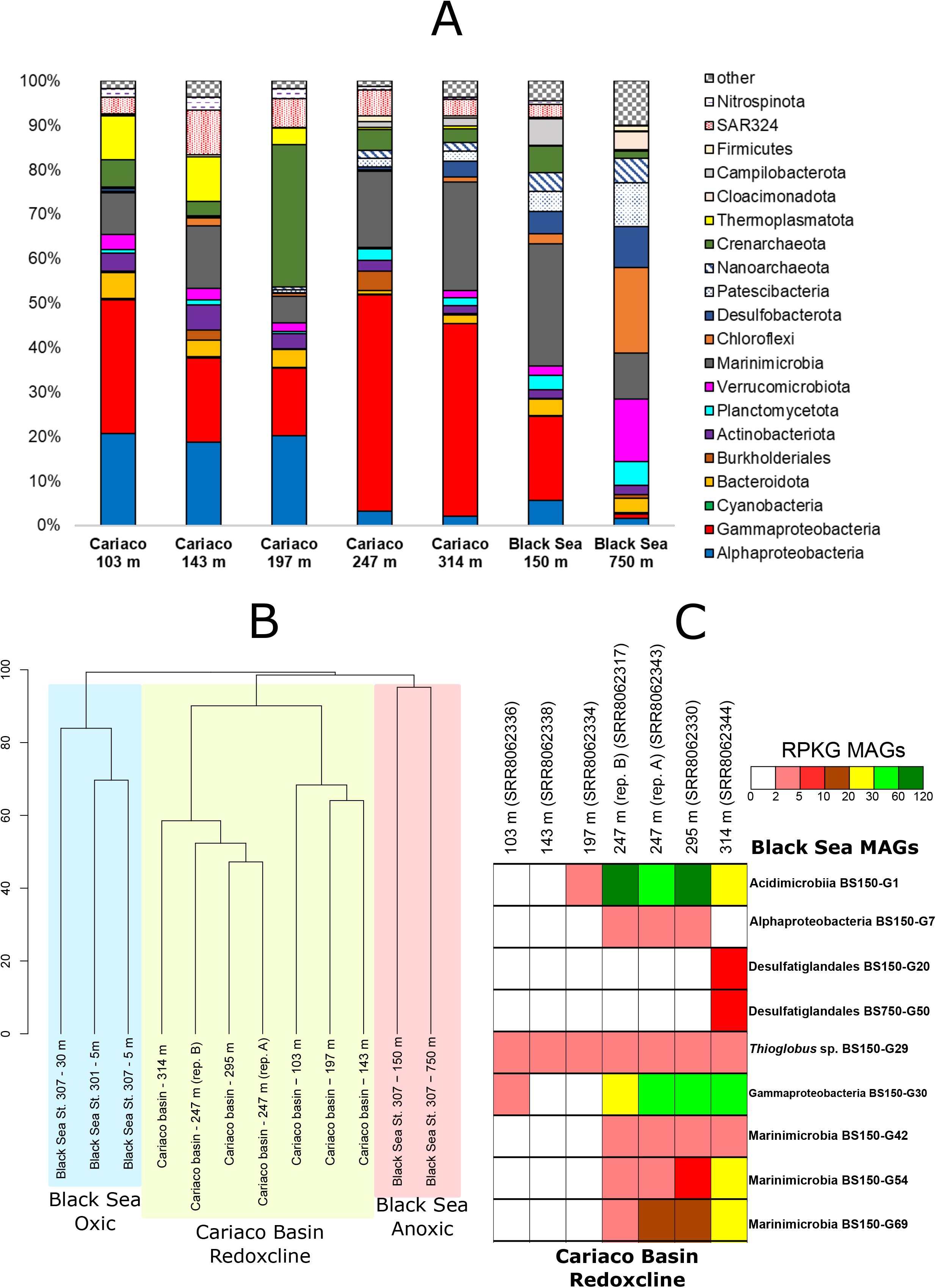
Comparison of Black Sea and Cariaco Basin redoxclines at the level of A) 16S rRNA taxonomic classification, B) Hierarchical read cluster analysis with Bray-Curtis presence/absence indexes and C) Black Sea species recruiting at the Cariaco depth profile datasets (PRJNA326482).

**Fig. S6.**
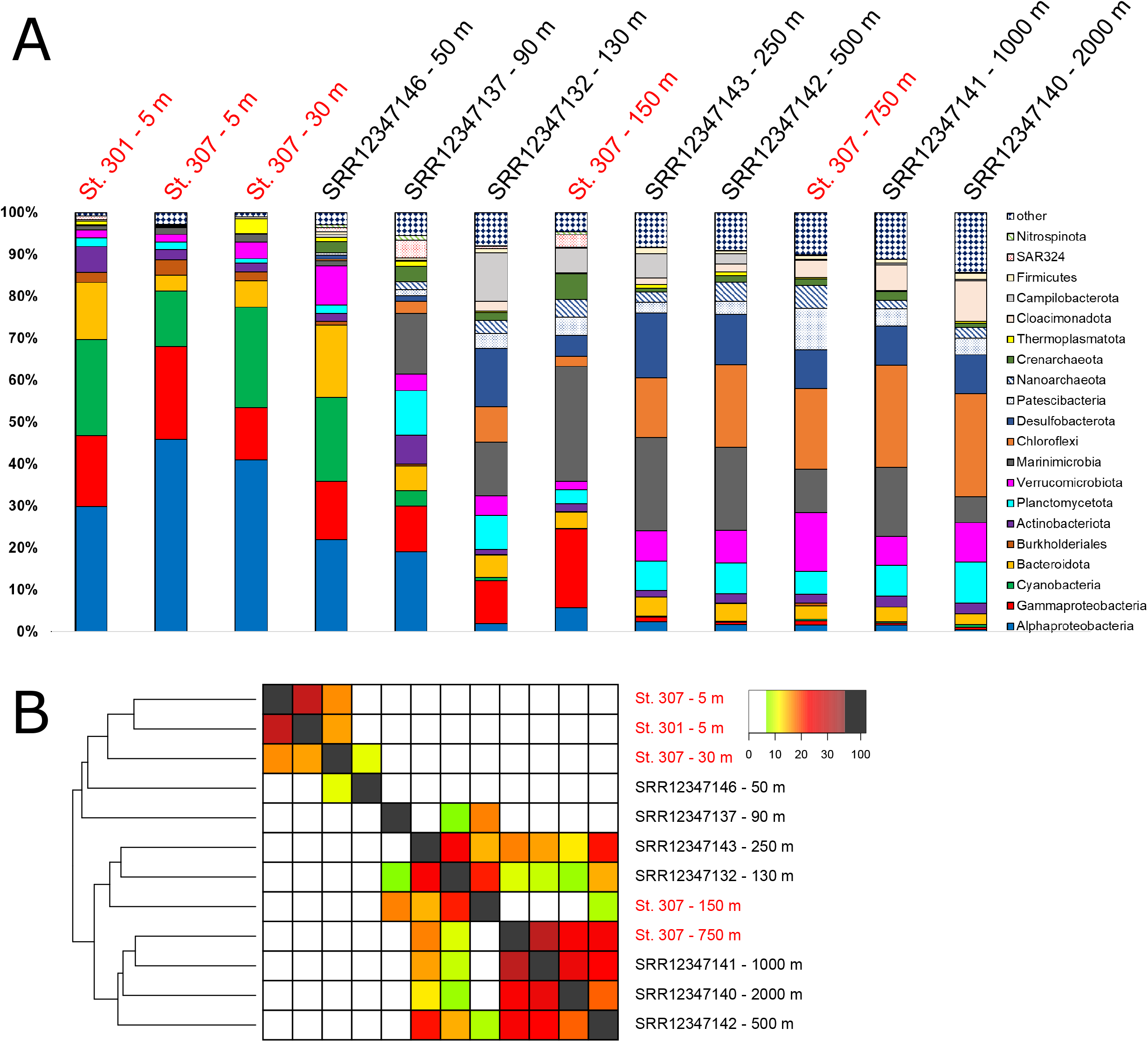
Comparison of Black Sea metagenomic datasets from the present study (in red) and those available from the NCBI database (Bioproject PRJNA649215). Comparison made at the level of A) Phylum 16S rRNA taxonomic classification, B) Heatmap read cluster analysis with Bray-Curtis presence/absence indexes.

## References

1. Konovalov SK, Murray JW, Luther III GW. Black Sea Biogeochemistry. Oceanography. 2005;18:24–35.

2. Stanev EV, He Y, Staneva J, Yakushev E. Mixing in the black sea detected from the temporal and spatial variability of oxygen and sulfide: Argo float observations and numerical modelling. European Geosciences Union; 2014;

3. Murray JW, Top Z, Özsoy E. Hydrographic properties and ventilation of the Black Sea. Deep Sea Res Part A Oceanogr Res Pap. Elsevier; 1991;38:S663–89.

4. Conley DJ, Bȷ̈orck S, Bonsdorff E, Carstensen J, Destouni G, Gustafsson BG, et al. Hypoxia-related processes in the Baltic Sea. Environ Sci Technol. ACS Publications; 2009;43:3412–20.

5. Suominen S, Dombrowski N, Sinninghe Damsté JS, Villanueva L. A diverse uncultivated microbial community is responsible for organic matter degradation in the Black Sea sulphidic zone. Environ Microbiol. Wiley Online Library; 2020;

6. Callieri C, Slabakova V, Dzhembekova N, Slabakova N, Peneva E, Cabello-Yeves PJ, et al. The mesopelagic anoxic Black Sea as an unexpected habitat for Synechococcus challenges our understanding of global “deep red fluorescence.” ISME J. Nature Publishing Group; 2019;1.

7. Di Cesare A, Dzhembekova N, Cabello-Yeves PJ, Eckert EM, Slabakova V, Slabakova N, et al. Genomic Comparison and Spatial Distribution of Different Synechococcus Phylotypes in the Black Sea. Front Microbiol. Frontiers; 2020;11:1979.

8. van Vliet DM, von Meijenfeldt FAB, Dutilh BE, Villanueva L, Sinninghe Damsté JS, Stams AJM, et al. The bacterial sulfur cycle in expanding dysoxic and euxinic marine waters. Environ Microbiol. Wiley Online Library;

9. Haro-Moreno JM, López-Pérez M, José R, Picazo A, Camacho A, Rodriguez-Valera F. Fine metagenomic profile of the Mediterranean stratified and mixed water columns revealed by assembly and recruitment. Microbiome. BioMed Central; 2018;6:128.

10. Mehrshad M, Amoozegar MA, Ghai R, Fazeli SAS, Rodriguez-Valera F. Genome reconstruction from metagenomic datasets reveals novel microbes in the brackish waters of the Caspian Sea. Appl Environ Microbiol. 2016;AEM. 03381–15.

11. Quast C, Pruesse E, Yilmaz P, Gerken J, Schweer T, Yarza P, et al. The SILVA ribosomal RNA gene database project: improved data processing and web-based tools. Nucleic Acids Res. Oxford University Press; 2012;41:D590–6.

12. Overbeek R, Olson R, Pusch GD, Olsen GJ, Davis JJ, Disz T, et al. The SEED and the Rapid Annotation of microbial genomes using Subsystems Technology (RAST). Nucleic Acids Res. 2013;42:D206–14.

13. Parks DH, Chuvochina M, Waite DW, Rinke C, Skarshewski A, Chaumeil P-A, et al. A standardized bacterial taxonomy based on genome phylogeny substantially revises the tree of life. Nat Biotechnol. Nature Publishing Group; 2018;

14. Cabello-Yeves PJ, Zemskaya TI, Zakharenko AS, Sakirko M V, Ivanov VG, Ghai R, et al. Microbiome of the deep Lake Baikal, a unique oxic bathypelagic habitat. Limnol Oceanogr. Wiley Online Library; 2019;

15. Scanlan DJ, Ostrowski M, Mazard S, Dufresne A, Garczarek L, Hess WR, et al. Ecological genomics of marine picocyanobacteria. Microbiol Mol Biol Rev. 2009;73:249–99.

16. Haro-Moreno JM, Rodriguez-Valera F, Rosselli R, Martinez-Hernandez F, Roda-Garcia JJ, Gomez ML, et al. Ecogenomics of the SAR11 clade. Environ Microbiol. Wiley Online Library; 2020;22:1748–63.

17. Hugerth LW, Larsson J, Alneberg J, Lindh M V, Legrand C, Pinhassi J, et al. Metagenome-assembled genomes uncover a global brackish microbiome. Genome Biol. 2015;16:1–18.

18. Overmann J, Cypionka H, Pfennig N. An extremely low-light adapted phototrophic sulfur bacterium from the Black Sea. Limnol Oceanogr. Wiley Online Library; 1992;37:150–5.

19. Marshall KT, Morris RM. Isolation of an aerobic sulfur oxidizer from the SUP05/Arctic96BD-19 clade. ISME J. Nature Publishing Group; 2013;7:452–5.

20. Callbeck CM, Lavik G, Ferdelman TG, Fuchs B, Gruber-Vodicka HR, Hach PF, et al. Oxygen minimum zone cryptic sulfur cycling sustained by offshore transport of key sulfur oxidizing bacteria. Nat Commun. Nature Publishing Group; 2018;9:1–11.

21. Suter EA, Pachiadaki M, Taylor GT, Astor Y, Edgcomb VP. Free-living chemoautotrophic and particle-attached heterotrophic prokaryotes dominate microbial assemblages along a pelagic redox gradient. Environ Microbiol. Wiley Online Library; 2018;20:693–712.

22. McLean JS, Bor B, Kerns KA, Liu Q, To TT, Solden L, et al. Acquisition and adaptation of ultra-small parasitic reduced genome bacteria to mammalian hosts. Cell Rep. Elsevier; 2020;32:107939.

23. Mara P, Vik D, Pachiadaki MG, Suter EA, Poulos B, Taylor GT, et al. Viral elements and their potential influence on microbial processes along the permanently stratified Cariaco Basin redoxcline. ISME J. Nature Publishing Group; 2020;1–14.

24. Marschall E, Jogler M, Henßge U, Overmann J. Large-scale distribution and activity patterns of an extremely low-light-adapted population of green sulfur bacteria in the Black Sea. Environ Microbiol. Wiley Online Library; 2010;12:1348–62.

25. Shah V, Zhao X, Lundeen RA, Ingalls AE, Nicastro D, Morris RM. Morphological plasticity in a sulfur-oxidizing marine bacterium from the SUP05 clade enhances dark carbon fixation. MBio. Am Soc Microbiol; 2019;10.

26. Bertagnolli AD, Padilla CC, Glass JB, Thamdrup B, Stewart FJ. Metabolic potential and in situ activity of marine Marinimicrobia bacteria in an anoxic water column. Environ Microbiol. Wiley Online Library; 2017;19:4392–416.

27. Hawley AK, Nobu MK, Wright JJ, Durno WE, Morgan-Lang C, Sage B, et al. Diverse Marinimicrobia bacteria may mediate coupled biogeochemical cycles along eco-thermodynamic gradients. Nat Commun. Nature Publishing Group; 2017;8:1–10.

28. Skennerton CT, Chourey K, Iyer R, Hettich RL, Tyson GW, Orphan VJ. Methane-fueled syntrophy through extracellular electron transfer: uncovering the genomic traits conserved within diverse bacterial partners of anaerobic methanotrophic archaea. MBio. Am Soc Microbiol; 2017;8:e00530–17.

29. Zhou Z, Pan J, Wang F, Gu J-D, Li M. Bathyarchaeota: globally distributed metabolic generalists in anoxic environments. FEMS Microbiol Rev. Oxford University Press; 2018;42:639–55.

30. Evans PN, Parks DH, Chadwick GL, Robbins SJ, Orphan VJ, Golding SD, et al. Methane metabolism in the archaeal phylum Bathyarchaeota revealed by genome-centric metagenomics. Science (80-). American Association for the Advancement of Science; 2015;350:434–8.

31. Laso-Pérez R, Wegener G, Knittel K, Widdel F, Harding KJ, Krukenberg V, et al. Thermophilic archaea activate butane via alkyl-coenzyme M formation. Nature. Nature Publishing Group; 2016;539:396–401.

32. Hamilton TL, Bovee RJ, Sattin SR, Mohr W, Gilhooly III WP, Lyons TW, et al. Carbon and sulfur cycling below the chemocline in a meromictic lake and the identification of a novel taxonomic lineage in the FCB superphylum, Candidatus Aegiribacteria. Front Microbiol. Frontiers; 2016;7:598.

33. Solli L, Håvelsrud OE, Horn SJ, Rike AG. A metagenomic study of the microbial communities in four parallel biogas reactors. Biotechnol Biofuels. Springer; 2014;7:146.

34. Anantharaman K, Brown CT, Hug LA, Sharon I, Castelle CJ, Probst AJ, et al. Thousands of microbial genomes shed light on interconnected biogeochemical processes in an aquifer system. Nat Commun. Nature Publishing Group; 2016;7:13219.

35. Nobu MK, Narihiro T, Rinke C, Kamagata Y, Tringe SG, Woyke T, et al. Microbial dark matter ecogenomics reveals complex synergistic networks in a methanogenic bioreactor. ISME J. Nature Publishing Group; 2015;9:1710–22.

36. Rinke C, Schwientek P, Sczyrba A, Ivanova NN, Anderson IJ, Cheng J-F, et al. Insights into the phylogeny and coding potential of microbial dark matter. Nature. Nature Publishing Group; 2013;499:431.

37. Castelle CJ, Brown CT, Anantharaman K, Probst AJ, Huang RH, Banfield JF. Biosynthetic capacity, metabolic variety and unusual biology in the CPR and DPANN radiations. Nat Rev Microbiol. Nature Publishing Group; 2018;16:629.

38. Li Y-X, Rao Y-Z, Qi Y-L, Qu Y-N, Chen Y-T, Jiao J-Y, et al. Deciphering symbiotic interactions of ‘Candidatus Aenigmarchaeota’with inferred horizontal gene transfers and co-occurrence networks. 2020;

39. Martín-Cuadrado A-B, López-García P, Alba J-C, Moreira D, Monticelli L, Strittmatter A, et al. Metagenomics of the deep Mediterranean, a warm bathypelagic habitat. PLoS One. 2007;2:e914.

40. Peng Y, Leung HCM, Yiu S-M, Chin FYL. IDBA-UD: a de novo assembler for single-cell and metagenomic sequencing data with highly uneven depth. Bioinformatics. 2012;28:1420–8.

41. Hyatt D, Chen G-L, LoCascio PF, Land ML, Larimer FW, Hauser LJ. Prodigal: prokaryotic gene recognition and translation initiation site identification. BMC Bioinformatics. 2010;11:1.

42. Buchfink B, Xie C, Huson DH. Fast and sensitive protein alignment using DIAMOND. Nat Methods. 2015;12:59–60.

43. Tatusov RL, Natale DA, Garkavtsev I V, Tatusova TA, Shankavaram UT, Rao BS, et al. The COG database: new developments in phylogenetic classification of proteins from complete genomes. Nucleic Acids Res. 2001;29:22–8.

44. Haft DH, Loftus BJ, Richardson DL, Yang F, Eisen JA, Paulsen IT, et al. TIGRFAMs: a protein family resource for the functional identification of proteins. Nucleic Acids Res. 2001;29:41–3.

45. Lowe TM, Eddy SR. tRNAscan-SE: a program for improved detection of transfer RNA genes in genomic sequence. Nucleic Acids Res. 1997;25:955–64.

46. Nawrocki EP, Eddy SR. ssu-align: a tool for structural alignment of SSU rRNA sequences. URL http://selab.janelia.org/software.html; 2010.

47. Kang D, Li F, Kirton ES, Thomas A, Egan RS, An H, et al. MetaBAT 2: an adaptive binning algorithm for robust and efficient genome reconstruction from metagenome assemblies. PeerJ Prepr. PeerJ Inc. San Diego, USA; 2019;7:e27522v1.

48. Lê S, Josse J, Husson F. FactoMineR: an R package for multivariate analysis. J Stat Softw. 2008;25:1–18.

49. Rice P, Longden I, Bleasby A. EMBOSS: the European molecular biology open software suite. Trends Genet. 2000;16:276–7.

50. Parks DH, Imelfort M, Skennerton CT, Hugenholtz P, Tyson GW. CheckM: assessing the quality of microbial genomes recovered from isolates, single cells, and metagenomes. Genome Res. 2015;25:1043–55.

51. Edgar RC. Search and clustering orders of magnitude faster than BLAST. Bioinformatics. 2010;26:2460–1.

52. Benoit G. Simka: fast kmer-based method for estimating the similarity between numerous metagenomic datasets. RCAM. 2015.

53. Legendre P, Anderson MJ. Distance-based redundancy analysis: testing multispecies responses in multifactorial ecological experiments. Ecol Monogr. Wiley Online Library; 1999;69:1–24.

54. Oksanen J, Kindt R, Legendre P, O’Hara B, Stevens MHH, Oksanen MJ, et al. The vegan package. Community Ecol Packag. 2007;10:631–7.

